# Piezo1 channel activates ADAM10 sheddase to regulate Notch1 and gene expression

**DOI:** 10.1101/732370

**Authors:** Vincenza Caolo, Marjolaine Debant, Naima Endesh, T Simon Futers, Laeticia Lichtenstein, Gregory Parsonage, Elizabeth AV Jones, David J Beech

## Abstract

Mechanical force has emerged as a determinant of Notch signalling but the mechanisms of force sensing and coupling to Notch are unclear. Here we propose a role for Piezo1 channels, the recently identified mechanosensors of mammalian systems. Piezo1 channel opening in response to shear stress or a chemical agonist led to activation of a disintegrin and metalloproteinase domain-containing protein 10 (ADAM10), a Ca^2+^-regulated transmembrane sheddase that mediates S2 Notch1 cleavage. Consistent with this observation there was increased Notch1 intracellular domain (NICD) that depended on ADAM10 and the downstream S3 cleavage enzyme, γ-secretase. Endothelial-specific disruption of Piezo1 in mice led to decreased Notch1-regulated gene expression in hepatic vasculature, consistent with prior evidence that Notch1 controls hepatic perfusion. The data suggest Piezo1 as a mechanism for coupling physiological force at the endothelium to ADAM10, Notch1, gene expression and vascular function.

## Introduction

Mammalian Notch proteins were identified following studies in *D. melanogaster* that linked genetic abnormality to wing notch^1^. Extensive research then revealed major roles in the transfer of information between cells in health and disease^1^. Each of the four Notch receptors (Notch1-4) is a membrane protein that is trans coupled to a membrane-anchored ligand such as Deltalike 4 (DLL4). Though the initiation of Notch signalling is often considered to occur through ligand-receptor complex formation, mechanical force also plays an important role in this activation whereby a pulling force arising from ligand endocytosis causes trans activation^1,2^. Furthermore it recently became apparent that frictional force from fluid flow also stimulates Notch1, but how this force couples to the Notch mechanism is unknown^3–6^. Therefore mechanical forces would seem to play key roles in Notch regulation. Further information is needed on how this is achieved.

Piezo1 channels are key players in the sensing of shear stress and lateral force applied to plasma membranes (membrane tension)^7–14^. While there are multiple candidate sensors, Piezo1 channels are notable because of broad agreement amongst investigators that they are direct sensors of physiological force. As such they are now considered to be bona fide force sensors rather than proteins that, whilst affected by force, do not appear to have evolved to sense and transduce force as a primary function. Piezo1 channels are exquisitely sensitive to membrane tension^15^ and readily able to confer force-sensing capacity on cells that are otherwise poorly sensitive^7,9^. Reconstitution of Piezo1 channels in artificial lipid bilayers generates force-sensing channels^16^ and native Piezo1 channels in excised membrane patches respond robustly to mechanical force in the absence of intracellular factors^10^.

Piezo1 channels are Ca^2+^-permeable non-selective cationic channels, so when force causes them to open there is Ca^2+^ entry, elevation of the cytosolic Ca^2+^ concentration and regulation of Ca^2+^-dependent mechanisms^7,8^. Potentially relevant to such a system is Ca^2+^ and Ca^2+^-calmodulin regulation of ADAM10^17,18^, a metalloprotease or sheddase that catalyses rate-limiting S2 cleavage of Notch1 prior to γ-secretase-mediated S3 cleavage and release of Notch1 intracellular domain (NICD), driving downstream transcription^1,19,20^. Therefore, we speculated about a potential relationship between Piezo1 and Notch1, using endothelial cells to investigate this idea because both proteins are prominent in these cells and have established functional significance in them^1,8–10,12,19^. Because mechanical force can affect numerous mechanisms, we explored the relationship by specifically activating Piezo1 channels with a synthetic small-molecule agonist (Yoda1) that acts directly to enhance force sensitivity^21–24^. Because there is inherent force in cell membranes and force arises through cell-cell and cell-substrate contact, Yoda1 can be used to activate the channels in endothelial cells without applying an exogenous force^24^ that might otherwise complicate the analysis by concurrently stimulating parallel mechanisms. Piezo1 dependence of Yoda1’s actions was investigated and the effect of fluid flow was also determined.

## Results

### Piezo1 regulates NICD abundance

To activate Piezo1 channels without stimulating other force-sensing mechanisms we applied 0.2 μM Yoda1, the concentration required for half-maximal activation of native endothelial Piezo1 channels^24^. After 30-min Yoda1 treatment, human microvascular endothelial cells (HMVEC-Cs) showed 1.5-2 times as much NICD as controls (Figure 1a-d). Short interfering (si) RNA specific to Piezo1 (Supplementary Figure S1) reduced basal NICD to about half its control value and the Yoda1 effect was suppressed (Figure 1b). The data are consistent with the reported ability of shear stress to increase NICD^4^ and suggest importance of Piezo1 for basal Notch1 processing and ability of Yoda1 to stimulate production of NICD.

**Figure 1:**
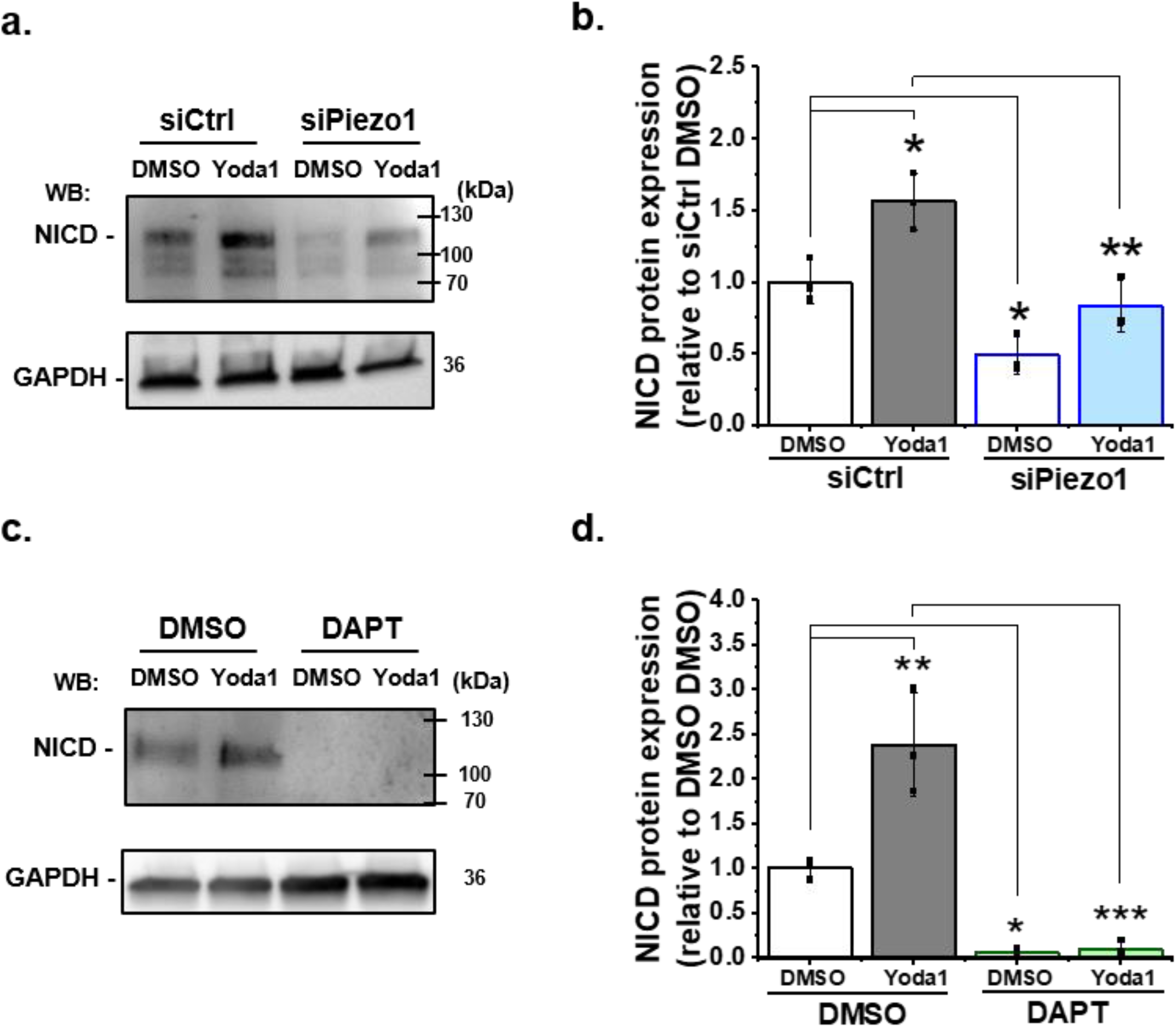
Piezo1 increases NICD abundance via γ-secretase. (**a**) Representative Western blot labelled with anti-NICD and anti-GAPDH (loading control) antibodies for HMVEC-Cs treated for 30 min with 0.2 µM Yoda1 or vehicle (DMSO) after transfection with control siRNA (siCtrl) or Piezo1 siRNA (siPiezo1). The expected mass of NICD is 110 kDa. Lower molecular bands were also apparent in some experiments and may have been degraded NICD. (**b**) Quantification of data of the type exemplified in (**a**), showing mean ± SD for abundance of NICD normalized to siCtrl DMSO (n = 3). (**c**) Representative Western blot labelled with anti-NICD and anti-GAPDH antibodies for HMVEC-Cs treated for 30 min with 0.2 µM Yoda1 or vehicle (DMSO) in the absence or presence of 10 µM DAPT. (**d**) Quantification of data of the type exemplified in (**c**), showing mean ± SD data for abundance of NICD normalized to vehicle (DMSO) control (n = 3). Statistical analysis: Two-way ANOVA test was used, indicating * P< 0.05, ** P< 0.01, *** P< 0.001.

### γ-secretase is required

If Piezo1 channels do indeed promote NICD release, the effect should depend on γ-secretase^1^. Therefore we tested the role of γ-secretase by treating cells with 10 μM DAPT (N-[N-(3,5-difluorophenacetyl)-l-alanyl]-S-phenylglycine t-butylester), a commonly used γ-secretase inhibitor^4,25^. DAPT had an effect on NICD that was similar to that of Piezo1 siRNA, reducing basal NICD and ablating the ability of Yoda1 to increase NICD (Figure 1c, d). DAPT inhibited Yoda1-evoked Ca^2+^ entry by about 30%, which was unlikely to have been sufficient to explain its effect on NICD (Supplementary Figure S1). The data suggest that Piezo1-mediated and constitutively generated NICD require γ-secretase.

### ADAM10 is important

In order to understand how Piezo1 channels couple to Notch1 we next considered the role of ADAM10, which is required for S2 cleavage before γ-secretase can act ^1^. SiRNA targeted to ADAM10 (Supplementary Figure S2) had an effect on NICD that was remarkably similar to that of Piezo1 siRNA and DAPT (Figure 2a, b *cf* Figure 1). It tended to reduce basal NICD, and Yoda1 no longer had significant effect (Figure 2b). The data suggest that ADAM10 is required for effects of Piezo1 on NICD.

**Figure 2:**
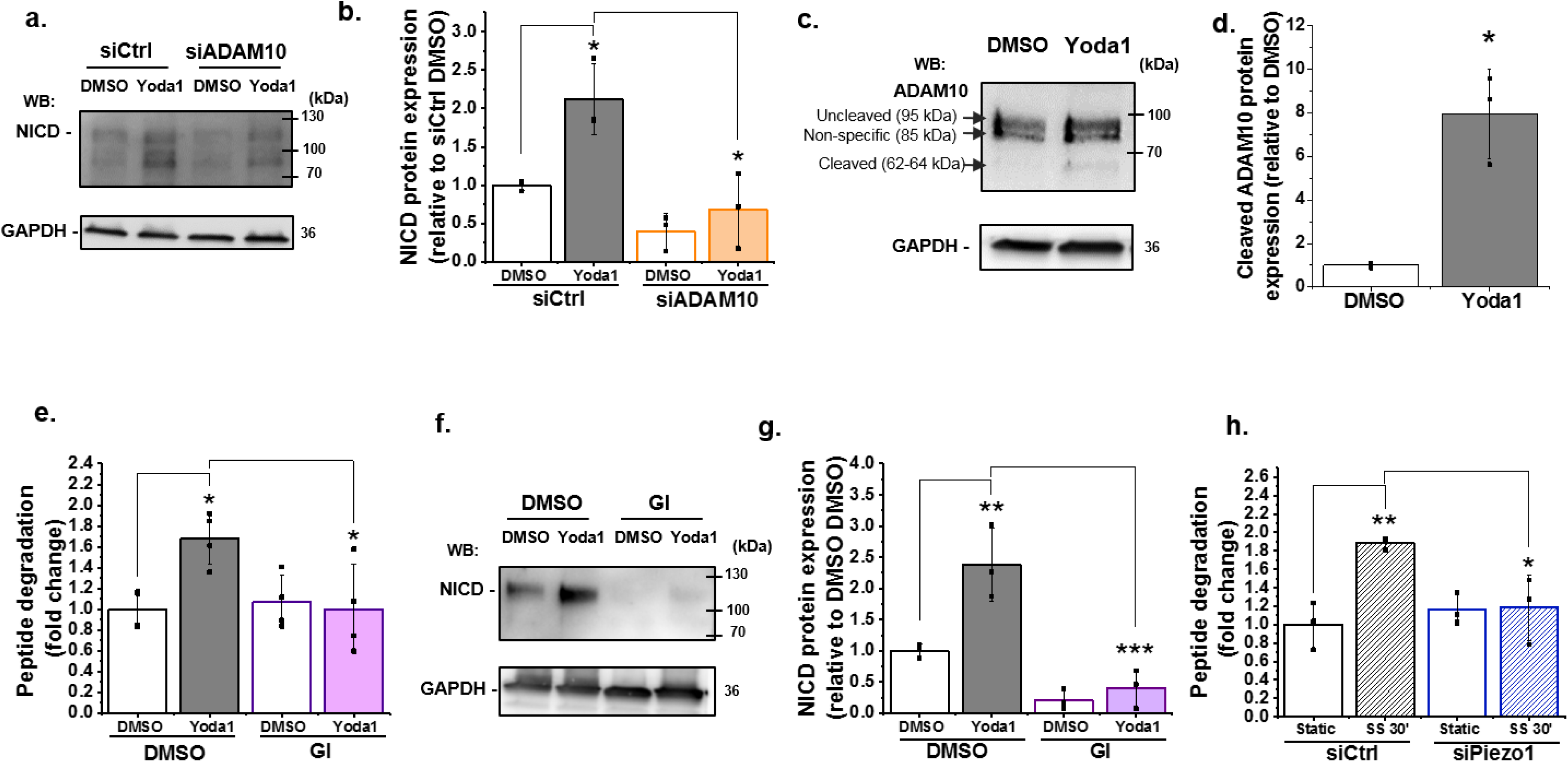
ADAM10 is important for Piezo1 regulation of NICD and activated by shear stress. (**a**) Representative Western blot labelled with anti-NICD and anti-GAPDH antibodies for HMVEC-Cs treated for 30 min with 0.2 µM Yoda1 or vehicle (DMSO) after transfection with control siRNA (siCtrl) or ADAM10 siRNA (siADAM10). (**b**) Quantification of data of the type exemplified in (**a**), showing mean ± SD data for abundance of NICD normalized to siCtrl DMSO (n = 3). (**c**, **d**) Quantification of uncleaved (95 kDa) and cleaved (62-64 kDa) ADAM10 in HMVEC-Cs after treatment for 30 min with Yoda1 (0.2 µM). The 85 kDa band between the uncleaved and cleaved ADAM10 was non-specific labelling not related to ADAM10 (Supplementary Figure S2). Data represent mean ± SD (n = 3) and normalization was to the reference protein, GAPDH. (**e**) ADAM10 enzyme activity assessed by specific peptide degradation and subsequent fluorescence emission after 30 min treatment of HMVEC-Cs with 0.2 µM Yoda1 in the absence or presence of 5 µM GI254023X (GI). Data are shown as mean ± SD data (n = 4) relative to DMSO condition. (**f**) Example Western blot labelled with anti-NICD and anti-GAPDH antibodies for HMVEC-Cs treated for 30 min with 0.2 µM Yoda1 or vehicle (DMSO) in the absence or presence of 5 µM GI254023X (GI). (**g**) Quantification of data of the type exemplified in (**f**), showing mean ± SD data for abundance of NICD normalized to DMSO (n = 3). (**h**) ADAM10 enzyme activity as in (**e**) but after 30 min exposure of HMVEC-Cs to 10 dyn.cm^−2^ laminar shear stress (SS). Static was without SS. Cells were transfected with control siRNA (siCtrl) or Piezo1 siRNA (siPiezo1). Data are shown as mean ± SD data (n = 3) relative to static condition. Statistical analysis: Two-way ANOVA test was used for (**b**, **e**, **g, h**), indicating * P< 0.05, ** P< 0.01, *** P< 0.001; t-Test was used for (**d**), indicating * P< 0.05, *** P< 0.001.

### ADAM10 enzyme activity is needed

To investigate if ADAM10 activity mediates the effect of Yoda1, we first quantified the abundance of its cleaved form, a 62-64 kDa protein that is generated by proprotein convertase to enable enzymatic activity^20^. In HMVEC-Cs, the majority of ADAM10 was in the uncleaved form, a protein of 95 kDa that is inactive (Figure 2c, Supplementary Figure S2). Yoda1 increased the abundance of the cleaved form, but the quantity remained small relative to the total protein (Figure 2c, d). To investigate if the ADAM10 was enzymatically active ^18^ we used a cell-based activity assay. Yoda1 increased ADAM10 activity (Figure 2e). The effect was prevented by the widely used ADAM10 inhibitor GI254023X ((2R,3S)-3-(Formyl-hydroxyamino)-2-(3-phenyl-1-propyl) butanoic acid[(1S)-2,2-dimethyl-1-methylcarbamoyl-1-propyl] amide) which is thought to act via the catalytic site^26^ (Figure 2e). GI254023X (5 μM) affected NICD similarly to Piezo1 siRNA and ADAM10 siRNA, reducing both basal NICD and the ability of Yoda1 to increase NICD (Figure 2f, g). A 10-fold lower concentration of GI254023X (500 nM) inhibited the Yoda1 effect on NICD (Supplementary Figure S2), consistent with its nanomolar potency against ADAM10^26^. GI254023X (5 μM) had no effect on Yoda1-evoked Ca^2+^ entry (Supplementary Figure S2). Gd^3+^ (30 μM), a blocker of the Piezo1 channel pore^7^, inhibited the ability of Yoda1 to activate ADAM10, consistent with Ca^2+^ entry through the channel being required (Supplementary Figure S2). The data suggest that Piezo1-mediated stimulation of ADAM10 enzyme activity is necessary for basal and stimulated effects on Notch1 cleavage.

### Shear stress activates ADAM10, via Piezo1

To further investigate ADAM10, we applied 10 dyn.cm^−2^ laminar shear stress, a physiological stimulator of Piezo1 channels^9^. In response to this stimulus there was also increased ADAM10 enzymatic activity, which was Piezo1-dependent (Figure 2h). The data suggest that shear stress acts, like Yoda1, to stimulate ADAM10 via Piezo1 channels.

### Function significance for downstream gene expression

To investigate if there is functional significance of Piezo1-regulated NICD, we quantified Notch1-regulated gene expression, focussing on the *HES1* gene which is Notch1- and flow-regulated^4^ and *DLL4*, which is itself Notch1 regulated^27^. Yoda1 caused increased expression of both *HES1* and *DLL4* genes (Figure 3), consistent with Yoda1’s effect on NICD (Figure 1). The effects of Yoda1 were suppressed by Piezo1 siRNA (Figure 3a, b), DAPT (Figure 3c, d), ADAM10 siRNA (Figure 3e, f) and GI254023X (Figure 3g, h). In most cases, there was little or no change in constitutive gene expression, suggesting that it is regulated mostly by other mechanisms in these cultured cells (HMVEC-Cs). Nevertheless, the data suggest that Piezo1 signalling via Notch1 can be functionally important for downstream gene regulation.

**Figure 3:**
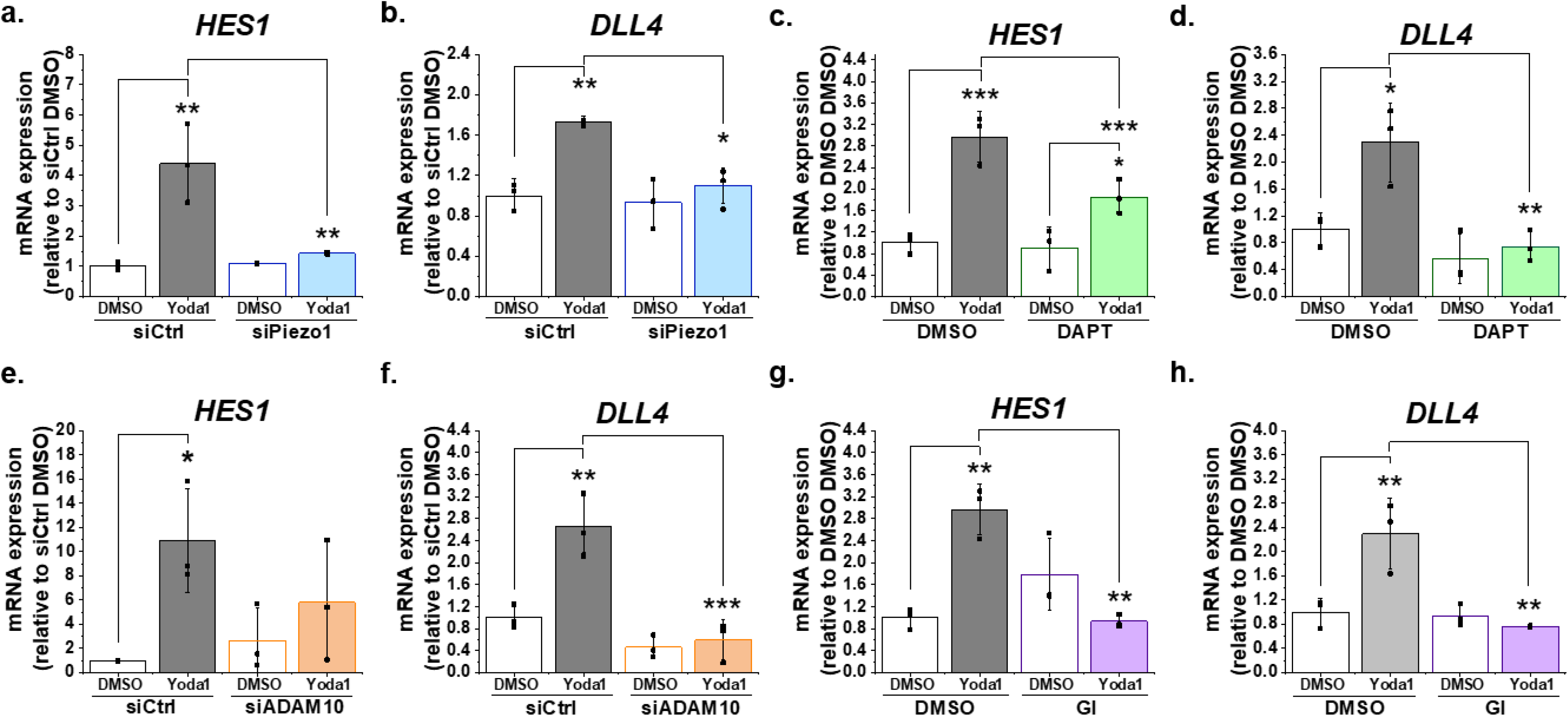
Function significance for downstream gene expression. (**a**, **b**) Summarized mean ± SD (n = 3) quantitative PCR data for fold-change in *HES1* (**a**) and *DLL4* (**b**) mRNA in HMVEC-Cs treated for 30 min with 0.2 µM Yoda1 or vehicle (DMSO) after transfection with control siRNA (siCtrl) or Piezo1 siRNA (siPiezo1). (**c**, **d**) Summarized mean ± SD (n = 3) quantitative PCR data for fold-change in *HES1* (**c**) and *DLL4* (**d**) mRNA in HMVEC-Cs treated for 30 min with 0.2 µM Yoda1 in the absence or presence of 10 µM DAPT for 2 hr. (**e**, **f**) Summarized mean ± SD (n = 3) quantitative PCR data for fold-change in *HES1* (**e**) and *DLL4* (**f**) mRNA in HMVEC-Cs treated for 30 min with 0.2 µM Yoda1 or vehicle (DMSO) after transfection with control siRNA (siCtrl) or ADAM10 siRNA (siADAM10). (**g**, **h**) Summarized mean ± SD (n = 3) quantitative PCR data for fold-change in *HES1* (**g**) and *DLL4* (**h**) mRNA in HMVEC-Cs treated for 30 min with 0.2 µM Yoda1 in the absence or presence of 5 µM GI254023X (GI) for 2 hr. Normalization and statistical analysis: mRNA expression was normalized to *GAPDH* mRNA abundance. Two-way ANOVA test was used, indicating * P< 0.05, ** P< 0.01, *** P< 0.001.

### Importance for hepatic vasculature gene regulation

To test the physiological relevance, we investigated liver where functional microvascular Piezo1 channels have been detected at the adult stage^10^ and where Notch1 is important for sinusoid development^19,28^. To specifically ask about endothelial biology, liver endothelial cells were acutely isolated after 2-weeks *in vivo* conditional Piezo1 disruption in endothelium of adult mice (Piezo1^ΔEC^), as previously described^10^. Neither mice nor isolated cells received Yoda1, so Piezo1 channel activity should have arisen only due to physiological forces existing in vivo prior to cell isolation. Strikingly, expression of *Hes1* and *Dll4* genes was downregulated in Piezo1^ΔEC^ endothelial cells (Figure 4a, b) (Supplementary Figure S3). *Hey1* and *EphrinB2*, other genes known to be Notch1- and flow-regulated^4,6^, were also downregulated (Figure 4c, d). Expression of the reference gene, *β-actin*, and endothelial marker gene, *Tie2*, was not affected (Supplementary Figure S3). Whole liver showed no change or trends towards decreased expression of *Hes1*, *Dll4*, *Hey1* and *Ephrin B2* in Piezo1^ΔEC^ mice, consistent with effects restricted to the endothelial cell population (Supplementary Figure S4). The data suggest that endothelial Piezo1 normally positively regulates Notch1 signalling in hepatic vasculature.

**Figure 4:**
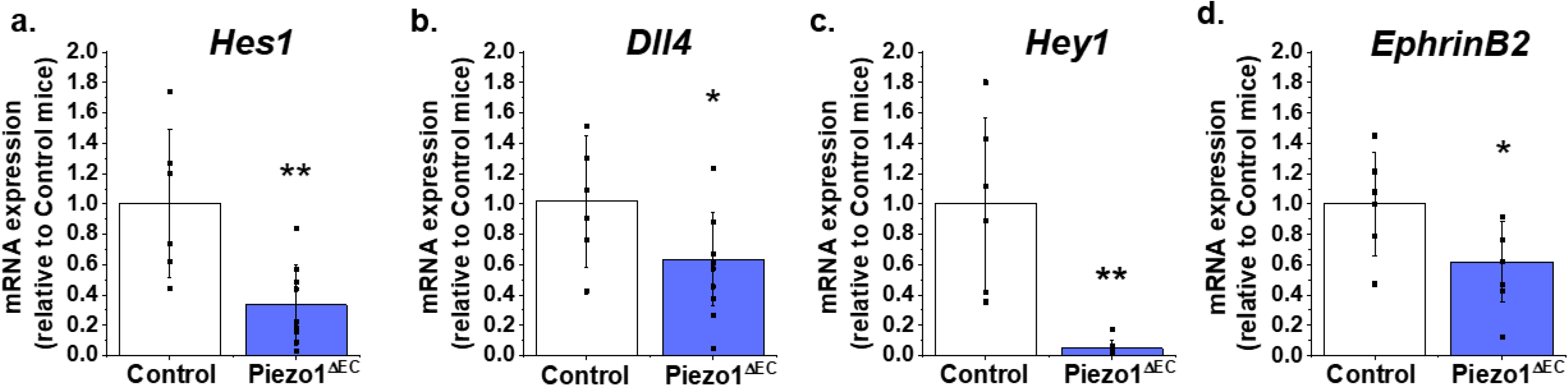
Endothelial Piezo1 promotes Notch1-regulated gene expression in liver. *Hes1* (**a**), *Dll4* (**b**), *Hey1* (**c**) and *EphrinB*2 (**d**) mRNA expression in liver endothelial cells freshly-isolated from control mice (Control, n = 6) and endothelial Piezo1 knockout mice (Piezo1^ΔEC^) (n = 9). Expression of all 4 genes was decreased in Piezo1^ΔEC^ compared with Control. Normalization and Statistical analysis: mRNA expression was normalized to abundance of β-actin mRNA, which was not different between Control and Piezo1^ΔEC^ (Supplementary Figure S3). t-Test was used, indicating significant difference of Piezo1^ΔEC^ *cf* Control * P< 0.05, ** P< 0.01.

## Discussion

The study has identified a connection between Piezo1 channels and Notch1 signalling and thus a mechanism by which Notch1 regulation by force can be achieved. We propose a pathway in which activation of Piezo1 channels stimulates ADAM10 via Ca^2+^ and Ca^2+^-calmodulin-regulated mechanisms^17,18^ for S2 cleavage of Notch1, which then enables intracellular S3 cleavage by γ-secretase and release of NICD for association with transcriptional regulators such as RBPJ and the control of gene expression (Figure 5). In this way, Notch1 is coupled to an exceptional force sensor, the Piezo1 channel. Other mechanisms by which Notch1 achieves force sensitivity are not excluded but we suggest that Piezo1 is a mechanism of biological significance because endothelial-specific disruption of Piezo1 in vivo disturbed Notch1-regulated gene expression in hepatic endothelium, a site known to be Notch-regulated. Our findings are consistent with prior studies of *D melanogaster* that inferred Notch to be downstream of Piezo ^29^ and so the proposed mechanism may have broad relevance.

**Figure 5:**
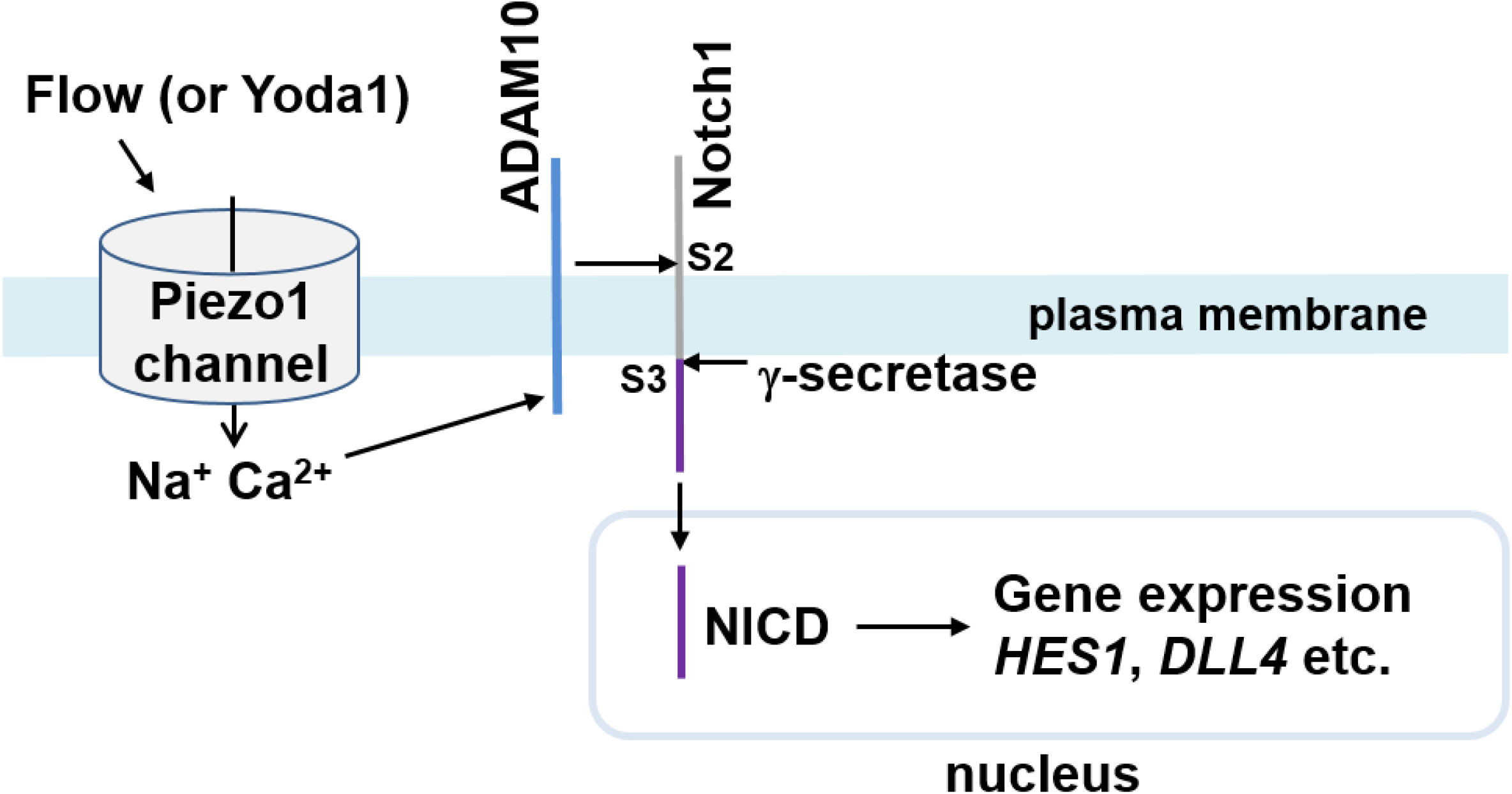
Summary of the proposed pathway. Activation of Piezo1 channel by mechanical force (e.g. fluid flow) or chemical agonist (Yoda1) causes elevation of the intracellular Ca^2+^ concentration which stimulates enzymatic activity of ADAM10 to cause S2 and S3 Notch1 cleavage and release of NICD to drive gene expression that is important in hepatic vasculature and potentially other vascular biology.

Prior work showed important effects of endothelial-specific disruption of Notch1-regulated RBPJ on hepatic microvasculature^19,28^. Disruption of RBPJ in 6 week-old mice led to enlarged sinusoids between portal and central venules, and disruption earlier in postnatal development caused poor perfusion, hypoxia and liver necrosis^28^. Therefore our observed requirement for Piezo1 in Notch1-regulated gene expression of hepatic endothelial cells suggests a positive role for the proposed Piezo1-ADAM10/Notch1 partnership in hepatic function. It is important to emphasise that Piezo1 signalling is unlikely to be limited to ADAM10/Notch1 because there is also good evidence for Piezo1 coupling to calpain^9^, eNOS^9,14^ and other mechanisms^30^. Therefore, it will be challenging to delineate the specific in vivo phenotypic contribution of Piezo1 coupling to ADAM10/Notch1 relative to other downstream consequences of activated Piezo1 channels.

An important observation of ours was that shear stress activates ADAM10 via Piezo1 (Figure 2h). It is consistent with previous work describing activation of ADAM10 by shear stress in human platelets^31^, which also express Piezo1^32^. The finding has implications for Notch1 signalling but also more widely because ADAM10 is a sheddase that also targets E- and VE-cadherin, CD44 and cellular prion protein^17,33–35^. Therefore its shear stress regulation via Piezo1 may be broadly relevant, for example in adherens junction biology and cartilage integrity, where functional importance of Piezo1 is already suggested^36,37^, and amyloid plaque formation, where Piezo1 was originally detected^38^ and has been suggested to have functional importance in combination with *E coli* infection^39^.

A prior report has suggested lack of specificity of Yoda1 for Piezo1 channels^40^ but the GsMTx4 toxin used as the basis of this proposal has limited value as a tool for evaluating Piezo1. Other prior studies have shown that genetic deletion of Piezo1 abolishes Yoda1’s effects^10,36,41,42^ and that Piezo1 knockdown by RNA interference suppresses its effects^14,30,43–46^, consistent with Yoda1 having only Piezo1-mediated effects. Specific structure-activity requirements of Yoda1 at Piezo1 have been observed^24^ and it does not activate Piezo2^21^. Here we observed that Piezo1-specific siRNA suppressed or abolished Yoda1 effects. Although it is important to seek Piezo1-specific activation with agents such as Yoda1 (because mechanical force is not specific to Piezo1) we recognise that activation by such agents does not necessarily mimic activation by a physiological factor such as shear stress. Importantly, therefore, we show that shear stress mimicked the effect of Yoda1 at ADAM10 and that genetic disruption of Piezo1 impacted Notch1 signalling in the absence of Yoda1. Therefore Piezo1 regulation of the ADAM10/Notch1 pathway was not dependent on Yoda1.

In conclusion, we connect the new discovery of Piezo1 mechanosensing with the extensive prior discoveries of the ADAM and Notch fields^1,19^. The ability of Piezo1 to activate ADAM10 and Notch1 suggests that these mechanisms are downstream of Piezo1 and therefore linked via Piezo1 to changes in physiological force. There are likely to be broad-ranging implications for vascular and other biology. Understanding how to specifically disrupt the Piezo1-ADAM10/Notch1 partnership will be important as a next step to deciphering the roles and therapeutic potential of the pathway.

## Materials and Methods

### Piezo1 mutant mice

All animal use was authorized by the University of Leeds Animal Ethics Committee and Home Office UK. Genotypes were determined using real-time PCR with specific probes designed for each gene (Transnetyx, Cordova, TN). C57BL/6J mice with *Piezo1* gene flanked with LoxP sites (Piezo1^flox^) were described previously^9^. To generate tamoxifen (TAM) inducible disruption of *Piezo1* gene in the endothelium (*Piezo1*^*ΔEC*^), Piezo1^flox^ mice were crossed with mice expressing cre recombinase under the Cadherin5 promoter (Tg(Cdh5-cre/ERT2)1Rha) and inbred to obtain Piezo1^flox/flox^/Cdh5-cre mice. TAM (T5648, Sigma-Aldrich, Saint-Louis, MO) was dissolved in corn oil (C8267 Sigma-Aldrich) at 20mg.ml^−1^. 10-12 week-old male mice were injected intra-peritoneal with 75 mg.kg^−1^ TAM for 5 consecutive days and studied 10-14 days later. Control mice were the same except they lacked cre, so they could not disrupt Piezo1 even though they were also injected with TAM.

### Acute isolation of liver endothelial cells

Liver of 12-14 week-old male mice was used. Tissue was mechanically separated using forceps, further cut in smaller pieces and incubated at 37 °C for 50 min, in a MACSMix Tube Rotator to provide continuous agitation, along with 0.1% Collagenase II (17101-015, Gibco, Waltham, MA) and Dispase Solution (17105-041, Gibco). Following enzymatic digestion samples were passed through 100 μm and 40 μm cell strainers to remove any undigested tissue. The suspension was incubated for 15 min with dead cell removal paramagnetic beads (130-090-101, Miltenyi Biotec GmbH, Bergisch Gladbach, Germany) and then passed through LS column (130-042-401, Miltenyi Biotec). The cell suspension was incubated with CD146 magnetic beads (130-092-007, Miltenyi Biotec 130-092-007) at 4°C for 15 min under continuous agitation and passed through MS column (130-042-201, Miltenyi Biotec). The CD146 positive cells, retained in the MS column, were plunge out with PEB and centrifuged at 1000 RPM for 5 min. Cell pellet was resuspended in RLT buffer (74004, Qiagen, Hilden, Germany) to proceed with RNA isolation.

### Cell culture

HMVEC-Cs were cultured in endothelial medium 2MV (EGM-2MV, CC-3202, Lonza, Basel, Switzerland) according to the manufacturer’s protocol. Sixteen hours before performing experiments, cells were cultured with starvation medium consisting of EGM-2MV but only 0.5% fetal bovine serum and without vascular endothelial growth factor A_165_ (VEGF A_165_) and basic fibroblast growth factor.

### siRNA transfection

HMVEC-Cs were transfected with siRNA using Opti-MEM^TM^ I Reduced Serum Medium (31985070, ThermoFisher Scientific, Waltham, MA) and Lipofectamine 2000 (11668019, ThermoFisher Scientific). For transfection of cells in 6-well plates, a total of 50 nmol siRNA in 0.1 mL was added to 0.8 mL cell culture medium per well. Medium was changed after 4 hours. After 48 hours cells were treated or not with Yoda1 and subjected to RNA or protein isolation. For Ca^2+^ measurement, cells were plated into a 96-well plate at a density of 25000 cells per well 24 hours after transfection, and Ca^2+^ entry was recorded 24 hours later.

### RNA isolation and RT-qPCR

For isolated liver endothelial cells, RNA was isolated by using RNeasy micro-kit (74004, Qiagen). A total of 100 ng RNA per sample was subjected to Reverse Transcriptase (RT) by using iScript^TM^ cDNA Synthesis kit (1708890, BioRad, Hercules, CA). For whole liver, RNA was isolated using phenol/chloroform extraction from snap frozen samples. One μg of RNA was used for RT (Superscript® III Reverse Transcriptase, 18080044, Invitrogen, Carlsbad, CA). qPCR was performed using SyBR Green (1725122, Biorad). The sequences of PCR primers are shown in Supplementary Table S1. Primers were synthetized by Sigma. qPCR reactions were performed on a LightCycler**®** 480 Real Time PCR System (Roche, Basel, Switzerland). Samples were analysed using the comparative CT method, where fold-change was calculated from the ΔΔCt values with the formula 2^−ΔΔCt^.

### ADAM10 enzyme activity

Activity was determined using the SensoLyte^®^520 ADAM10 Activity Assay Kit (AS-72226, AnaSpec Inc, Fremont, CA), which is based on the FRET substrate 5‐ FAM/QXL^™^520 with excitation/emission of 490/520 nm. HMVEC-Cs were treated with or without Yoda1 for 30 min in the presence or absence of ADAM10 inhibitor. Cells were then washed with PBS and collected with Trypsin-EDTA. The pellet was resuspended in assay buffer, incubated on ice for 10 min and centrifuged at 10 000 *g* for 10 min at 4°C. The supernatants were plated on a 96-well plate. The substrate solution was diluted in Assay Buffer, brought to 37 °C was then mixed 1:1 with the sample. The fluorescence was measured every 2.5 min for 60 min at 37 °C at excitation/emission of 490/520 nm Flexstation 3 microplate reader with SoftMax Pro 5.4.5 software (Molecular Devices, San Josa, CA).

### Shear stress

Endothelial cells were seeded on glass slides (MENSJ5800AMNZ, VWR, Radnor, PA) coated with Fibronectin (F0895, Sigma-Aldrich). Sixteen hours before performing experiments, cells were cultured with starvation medium consisting of EGM-2MV but only 0.5% fetal bovine serum and without vascular endothelial growth factor A_165_ (VEGF A_165_) and basic fibroblast growth factor. The slides were placed in a parallel flow chamber and flow of starvation medium was driven using a peristaltic pump.

### Measurement of intracellular Ca^2+^ concentration ([Ca^2+^]_i_)

Cells plated in 96-well plates were incubated for 1 hour in Standard Bath Solution (SBS, containing in mM: 130 NaCl, 5 KCl, 8 D-glucose, 10 HEPES, 1.2 MgCl_2_, 1.5 CaCl_2_, pH 7.4) supplemented with 2 μM fura-2-AM (F1201, Molecular Probes, Eugene, OR) and 0.01 % pluronic acid. Cells were then washed in SBS at room temperature for 30 min, allowing deesterification to release free fura-2. Fluorescence (F) acquisition (excitation 340 and 380 nm; emission 510 nm) was performed on a Flexstation 3 microplate reader with SoftMax Pro 5.4.5 software (Molecular Devices). After 60 seconds of recording, Yoda1 was injected. Ca^2+^ entry was quantified after normalization (ΔF340/380 = F340/380(t)-F340/380(t=0)).

### Immunoblotting

Proteins were isolated in RIPA buffer supplemented with PMSF, protease inhibitor mixture, and sodium orthovanadate (RIPA Lysis Buffer System, sc24948, Santa Cruz, Dallas, TX). Samples were heated at 95 °C for 5 min in SDS-PAGE sample buffer, loaded on a precast 4-20 % polyacrylamide gradient gel (4561094, Biorad) and subjected to electrophoresis. Proteins were transferred onto a nitrocellulose membrane (Trans-Blot^**®**^ Turbo^TM^ RTA Mini Nitrocellulose Transfer Kit, 1704270, BioRad) for 30 min using Trans-Blot Turbo Transfer System (BioRad). Membranes were blocked with 5 % milk in Tris-buffered saline with Tween 0.05 % for 1 hour at room temperature. The membranes were exposed to primary antibody overnight at 4 °C, rinsed and incubated with appropriate horseradish peroxidase-labelled secondary antibody for 1 hour at room temperature. The detection was performed by using SuperSignal^TM^ West Femto (34096, ThermoFisher Scientific) and visualized with a G-Box Chemi-XT4 (SynGene, Cambridge, UK). GAPDH was used as reference protein.

### Reagents

Human cardiac microvascular endothelial cells (HMVEC-C, CC-7030, Lonza), DAPT (D5942, Sigma-Aldrich), GI254023X (3995, Tocris Bioscience, Bristol, UK), Yoda1 (5586/10, Tocris Bioscience), ON-TARGET plus Control siRNA (Dharmacon, Lafayette, CO), siRNA Piezo1 (Sigma-Aldrich: 5’-GCAAGUUCGUGCGCGGAUU[dT][dT]-3’), ON-TARGET plus SMARTpool human siRNA ADAM10 (Dharmacon), cleaved Notch1 Val1744 D3B8 rabbit monoclonal (4147, Cell Signaling Technology, Danvers, MA), rabbit anti-ADAM10 (AB19026, Merck KGaA, Darmstadt, Germany), goat anti human VEGFR2 (AF357, R&D system, Minneapolis, MN), mouse anti-human PECAM-1 (CD31) (M0823, Agilent Dako, Santa Clara, CA), GAPDH mouse anti-human (10R-G109b, Fitzgerald Industries International, Acton, MA) and anti-mouse, anti-rabbit and anti-goat HRP conjugated secondary antibodies (Jackson ImmunoResearch, Ely, UK).

### Statistical Analysis

All averaged data are presented as mean ± standard deviation (SD). Statistical significance was determined using two-tailed *t*-test when only two groups were compared or by 2-way ANOVA followed by Tukey posthoc test when multiple groups were treated with vehicle control (DMSO) or Yoda1 were studied. The genotypes of mice were blinded to the experimental investigator and studied at random according to Mendelian ratio. In all cases, statistical significance was assumed for probability (*P*) <0.05. Statistical tests were performed using OriginPro 8.6 software or GraphPad Prism 6.0. Outlying data were not detected or excluded. The letter n indicates the number of independent biological experiments and its value in each case is stated in figure legends. The study was aimed at discovering components of a biological mechanism using various cell/molecular and animal-based studies to address a single hypothesis. In the absence of prior knowledge of the mechanism, power calculations were not considered to be applicable. We selected numbers of independent repeats of experiments base on prior experience of studies of this type. In all cases the number of independent repeats was at least 3. The number of replicates per independent experiment was 1 for western blotting, 2 for qPCR, 4 for Ca^2+^ assays and 1 for the ADAM10 activity assay.

## Supplementary Information

Supplementary information is provided as a separate document. Included in this document are images of the uncropped western blots presented concisely in the main figures (Supplementary Figure S5).

## Acknowledgements

The study was supported by EU Marie Skłodowska Curie Individual Fellowship to VC and by grants from the Wellcome Trust and British Heart Foundation to DJB and a Fonds Wetenschappelijk Onderzoek grant G091018N to EAVJ. We thank Fiona Bartoli for PCR primer design and Richard Cubbon for helpful comments on the manuscript.

## Supplementary Information

**Supplementary Table S1.**
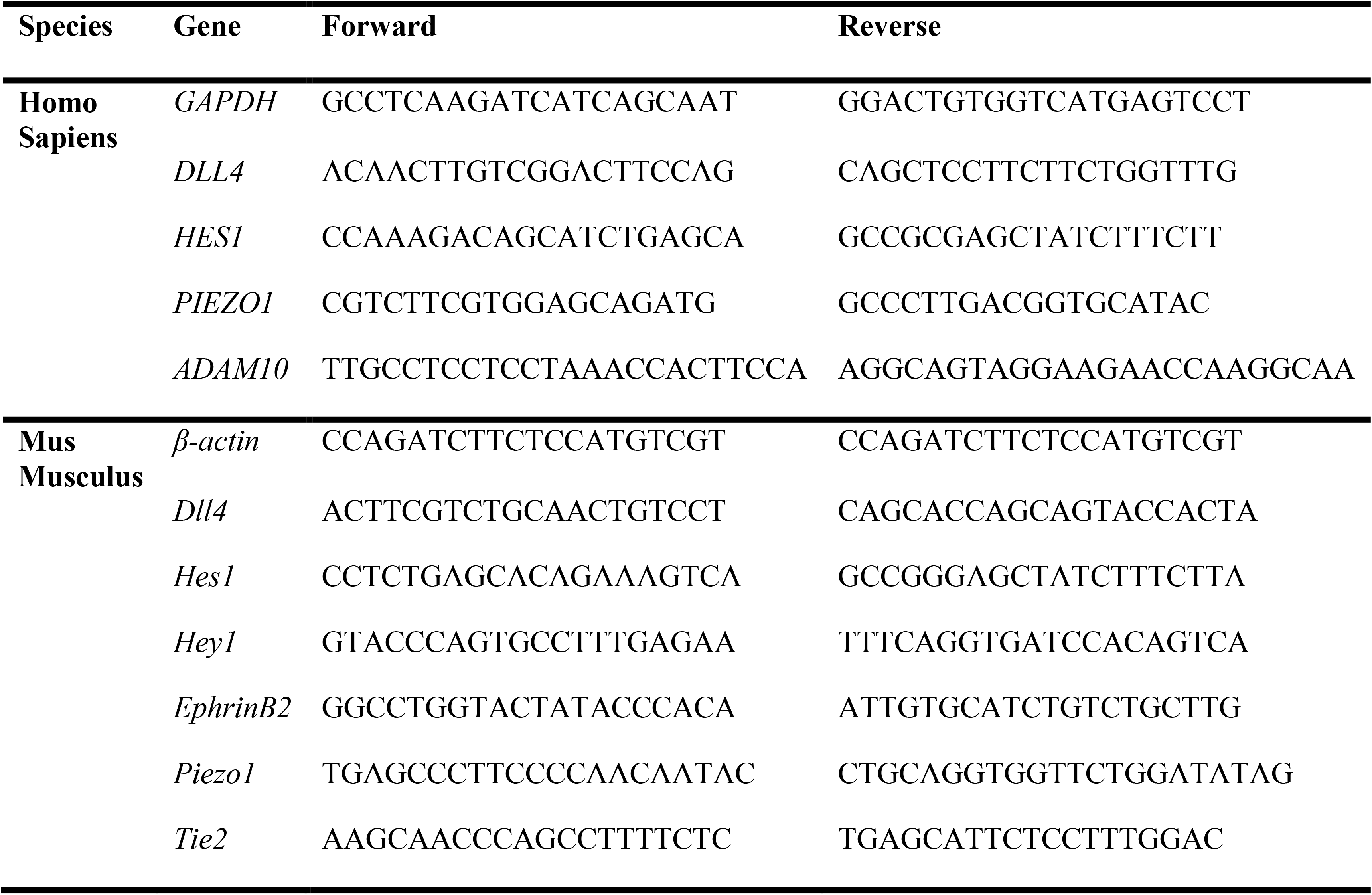
PCR primer sequences.

**Supplementary Figure S1:**
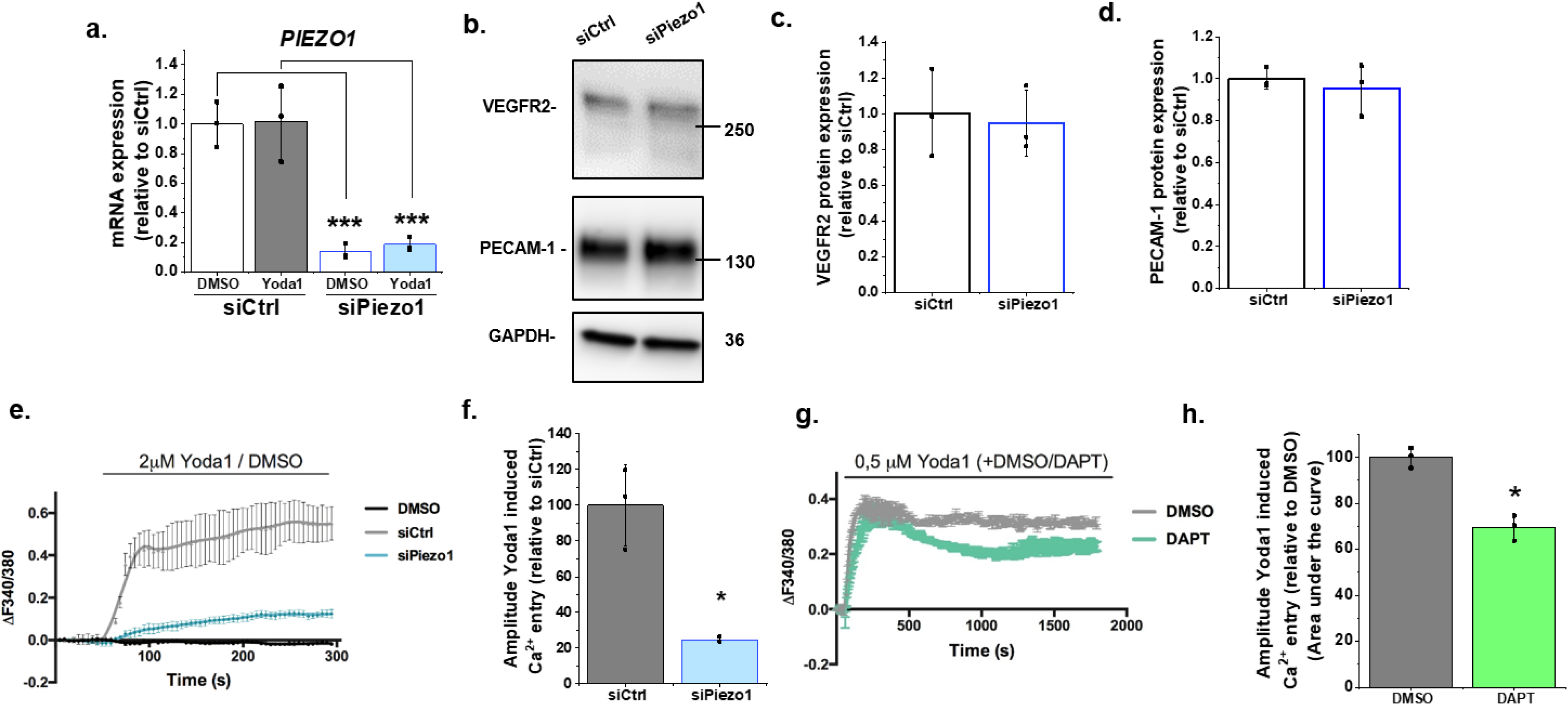
Supporting data for main Figure 1. (**a**) Summarized mean ± SD (n = 3) quantitative PCR data for fold-change in Piezo1 mRNA in HMVEC-Cs treated for 30 min with 0.2 μM Yoda1 or vehicle (DMSO) after transfection with control siRNA (siCtrl) or Piezo1 siRNA (siPiezo1). **(b)** Representative Western blot labelled with anti-VEGRF2, anti-PECAM-1 (anti-CD31) and anti-GAPDH antibodies for HMVEC-Cs after transfection with control siRNA (siCtrl) or Piezo1 siRNA (siPiezo1). (**c, d**) Quantification of data of the type exemplified in (**b**), showing mean ± SD data for abundance of VEGFR2 (**c**) and PECAM-1 (**d**) normalized to siCtrl (n = 3). (**e-f**) Representative intracellular Ca^2+^ measurement traces (**e**) in HMVEC-Cs during application of 2 μM Yoda1 or its control DMSO, 48 hours after transfection with control siRNA (siCtrl) or Piezo1 siRNA (siPiezo1), shown as mean ± SD of the amplitude for n = 3 (**f**). (**g-h**) Representative intracellular Ca^2+^ measurement traces (**g**) in HMVEC-Cs during prolonged application of 0.5 μM Yoda1 or its vehicle control (DMSO) simultaneously with 10 μM DAPT or its vehicle control (DMSO), shown as mean ± SD of the area under the curve for n = 3 (**h**). Two-way ANOVA test was used for (**a**) and t-test for (**c**, **d**, **f**, **h**), indicating * P< 0.05, *** P< 0.001.

**Supplementary Figure S2:**
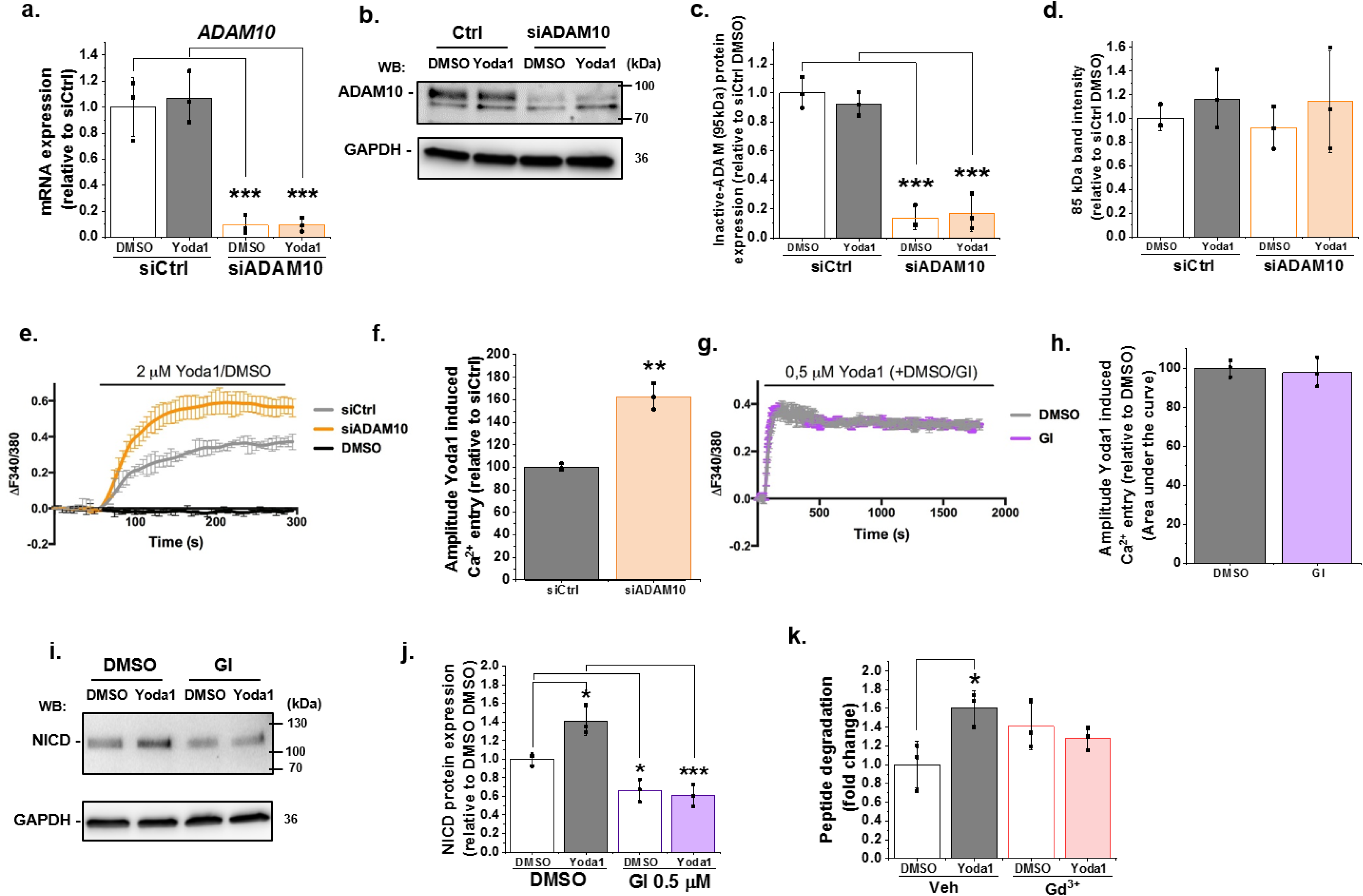
Supporting data for main Figure 2. (**a**) Summarized mean ± SD (n = 3) quantitative PCR data for fold-change in ADAM10 mRNA in HMVEC-Cs treated for 30 min with 0.2 µM Yoda1 or vehicle (DMSO) after transfection with control siRNA (siCtrl) or ADAM10 siRNA (siADAM10). **(b)** Representative Western blot labelled with anti-ADAM10 antibody for HMVEC-Cs after transfection with control siRNA (siCtrl) or ADAM10 siRNA (siADAM10). (**c, d**) Quantification of data of the type exemplified in (**b**), showing mean ± SD data for abundance of uncleaved ADAM10 (95 kDa) (**c**) and non-specific labelling of an unknown protein (85 kDa) (**d**) normalized to siCtrl (n = 3). (**e-f**) Representative intracellular Ca^2+^ measurement traces (**e**) in HMVEC-Cs during application of 2 μM Yoda1 or its vehicle control (DMSO), 48 hours after transfection with control siRNA (siCtrl) or ADAM10 siRNA (siADAM10), shown as mean ± SD of the amplitude for n = 3 (**f**). (**g-h**) Representative intracellular Ca^2+^ measurement traces (**g**) in HMVEC-Cs during prolonged application of 0.5 μM Yoda1 or its vehicle control (DMSO) simultaneously with 5 µM GI254023X (GI) or its vehicle control (DMSO), shown as mean ± SD of the area under the curve for n = 3 (**h**). (**i**) Representative Western blot labelled with anti-NICD and anti-GAPDH antibodies for HMVEC-Cs treated for 30 min with 0.2 µM Yoda1 or vehicle (DMSO) in the absence or presence of 0.5 µM GI254023X (GI) (**j**) Quantification of data of the type exemplified in (**i**), showing mean ± SD data for abundance of NICD normalized to siCtrl DMSO (n = 3). **(k)** ADAM10 enzyme activity assessed by specific peptide degradation and subsequent fluorescence emission after 30 min treatment of HMVEC-Cs with 0.2 µM Yoda1 in the absence or presence of 30 µM Gd^3+^. Data are shown as mean ± SD data (n = 3) relative to vehicle condition. Statistical analysis: Two-way ANOVA test was used, indicating * P< 0.05, ** P< 0.01 and *** P< 0.001

**Supplementary Figure S3:**
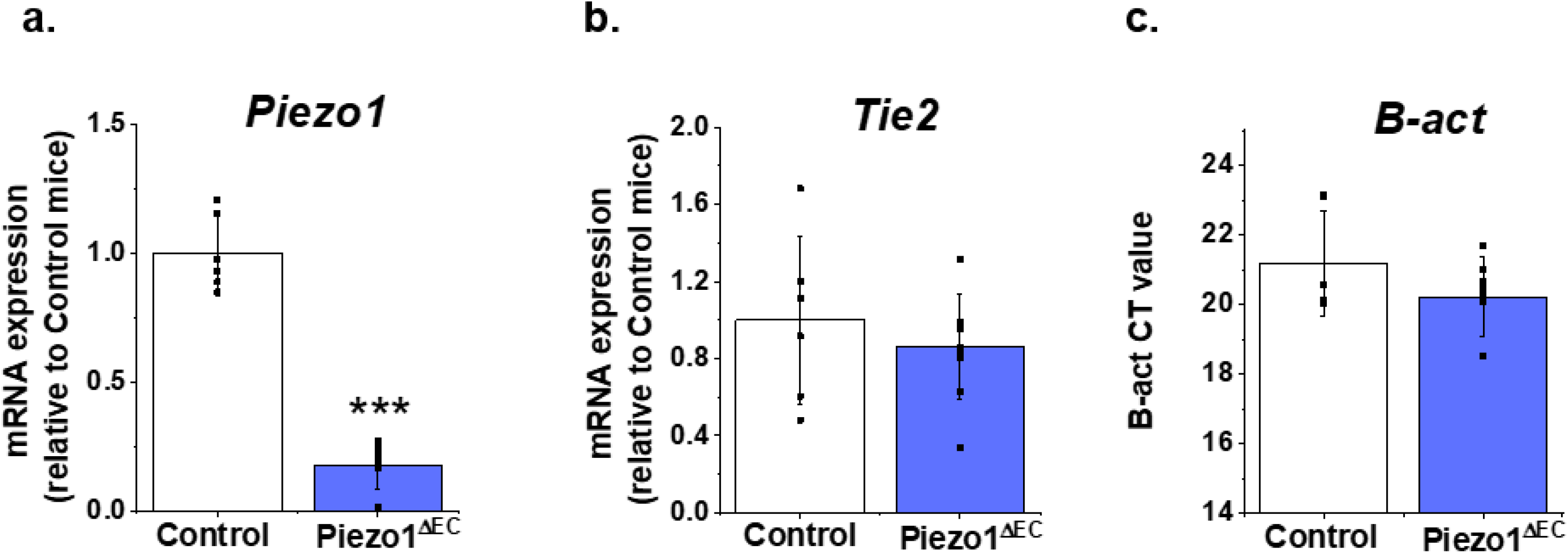
Supporting data for main Figure 4. Additional gene quantification for liver endothelial cells from Piezo1^ΔEC^ and Control mice. (**a**) *Piezo1*. (**b**) *Tie2*. (**c**) *β-actin* (raw CT values). (**a**, **b**) Normalized to abundance of *β-actin* (reference) expression. Statistical analysis: t-test was used for comparisons, *** P< 0.001.

**Supplementary Figure S4:**
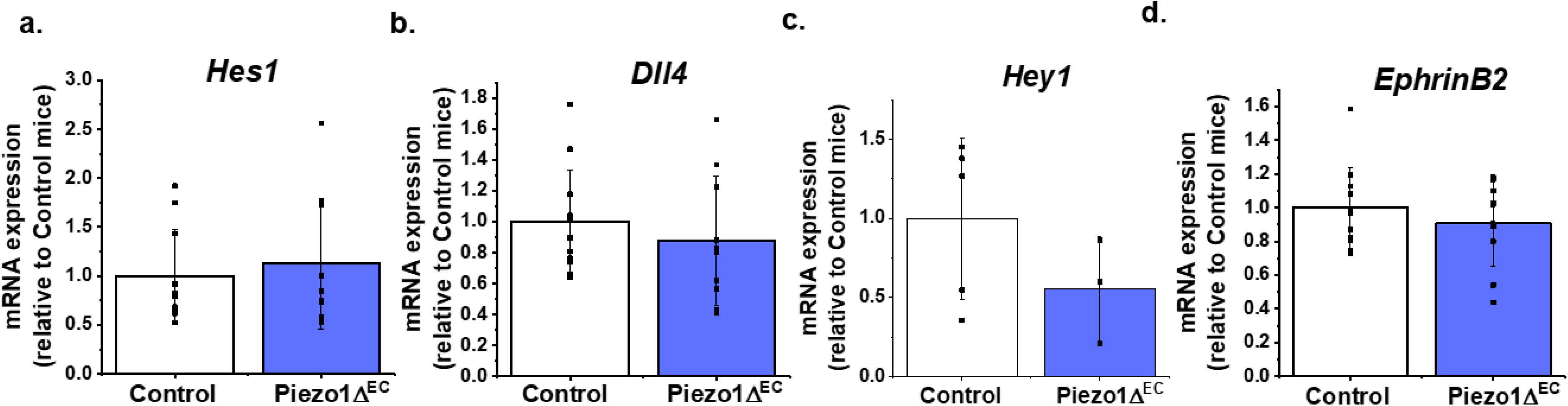
Notch1-regulated gene expression in whole liver. Gene quantification for whole liver from Piezo1^ΔEC^ (n=10) and Control mice (n=12). (**a**) *Hes1*. (**b**) *Dll4*. (**c**) *Hey1* (in 7 Control and 7 Piezo1^ΔEC^ experiments *Hey1* expression was not convincingly detected, so these data were excluded). (**d**) *Ephrin B2*. Normalized to abundance of the reference *β-actin* expression. Statistical analysis: t-test was used for comparisons. There were no significant differences between any pairs.

**Supplementary Figure S5:**
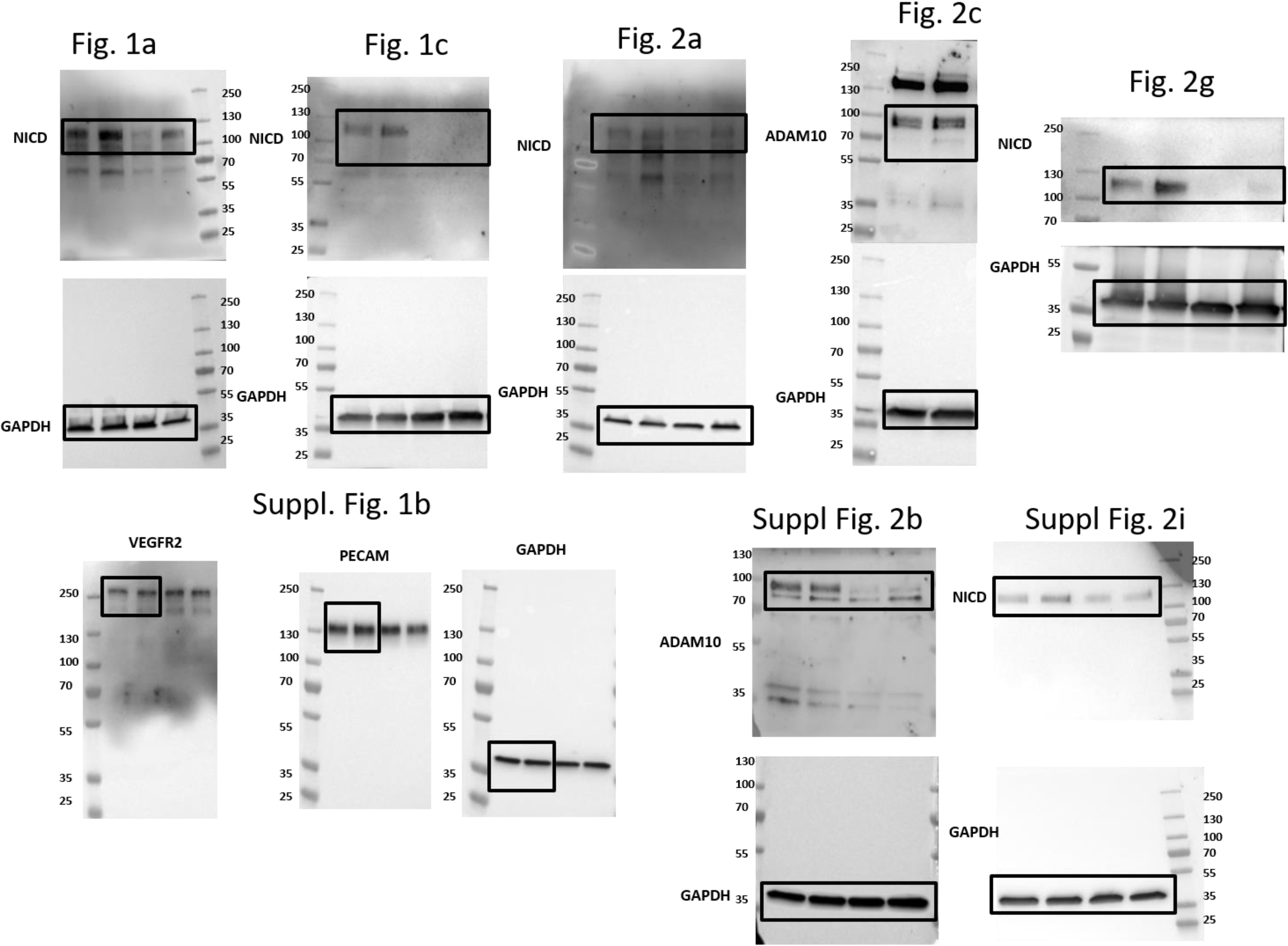
Uncropped western blots.

